# Lipoarabinomannan regulates septation in *Mycobacterium smegmatis*

**DOI:** 10.1101/2023.03.26.534150

**Authors:** Ian L. Sparks, Japinder Nijjer, Jing Yan, Yasu S. Morita

## Abstract

The growth and division of mycobacteria, which include several clinically relevant pathogens, deviate significantly from that of canonical bacterial models. Despite their Gram-positive ancestry, mycobacteria synthesize and elongate a diderm envelope asymmetrically from the poles, with the old pole elongating more robustly than the new pole. In addition to being structurally distinct, the molecular components of the mycobacterial envelope are also evolutionarily unique, including the phosphatidylinositol-anchored lipoglycans lipomannan (LM) and lipoarabinomannan (LAM). LM and LAM modulate host immunity during infection, but their role outside of intracellular survival remains poorly understood, despite their widespread conservation among non-pathogenic and opportunistically pathogenic mycobacteria. Previously, *Mycobacterium smegmatis* and *Mycobacterium tuberculosis* mutants producing structurally altered LM and LAM were shown to grow slowly under certain conditions and to be more sensitive to antibiotics, suggesting that mycobacterial lipoglycans may support cellular integrity or growth. To test this, we constructed multiple biosynthetic lipoglycan mutants of *M. smegmatis* and determined the effect of each mutation on cell wall biosynthesis, envelope integrity, and division. We found that mutants deficient in LAM, but not LM, fail to maintain cell wall integrity in a medium-dependent manner, with envelope deformations specifically associated with septa and new poles. Conversely, a mutant producing abnormally large LAM formed multiseptated cells in way distinct from that observed in a septal hydrolase mutant. These results show that LAM plays critical and distinct roles at subcellular locations associated with division in mycobacteria, including maintenance of local cell envelope integrity and septal placement.

**Significance:** Mycobacteria cause many diseases including tuberculosis (TB). Lipoarabinomannan (LAM) is a lipoglycan of mycobacteria and related bacteria, playing important roles as a surface-exposed pathogen-associated molecular pattern during host-pathogen interactions. Its importance is highlighted by the facts that anti-LAM antibody appears to be protective against TB disease progression, and urine LAM serves as a diagnostic marker for active TB. Given the clinical and immunological relevance of the molecule, it was a striking gap in knowledge that we did not know the cellular function of this lipoglycan in mycobacteria. In this study, we demonstrated that LAM regulates septation, a principle potentially generalizable to other lipoglycans widely found in a group of Gram-positive bacteria that lack lipoteichoic acids.

## Introduction

Cell envelopes define the shape, growth characteristics, and environmental interactions of bacteria and are broadly classified as Gram-positive or Gram-negative based on their genetically encoded envelope architecture. *Mycobacterium*, a medically important genus of actinobacteria, produces an envelope divergent from both of these regimes and consequently have drastically different envelope physiology. The mechanisms that govern the cell envelope assembly are only beginning to emerge.

Like other bacteria, mycobacteria have a plasma membrane and a peptidoglycan cell wall. However, mycobacteria also have an arabinogalactan layer covalently bound to their peptidoglycan, which is also bound to long chain fatty acids called mycolic acids (1–4). These bound mycolic acids support an outer “myco” membrane composed of free mycolic acids and other lipids. Altogether, this envelope architecture is lipid-dense, and includes multiple hydrophobic and hydrophilic layers, making it difficult for antibiotics to reach their targets. The structural divergence of mycobacterial envelopes also extends to the mechanisms used by mycobacteria to synthesize these structures during growth and division. Mycobacteria grow asymmetrically from the poles, with the old pole elongating faster than the new pole formed from the most recent cell septation event (5–8). While we have a basic understanding of the order of events during growth and cell division as well as genes that are involved in the biosynthesis of each component, many of the structural components of the cell envelope have poorly understood physiological functions. Examples include the phosphatidylinositol mannosides (PIMs) and their derivative lipoglycans lipomannan (LM) and lipoarabinomannan (LAM), which are phosphatidylinositol (PI)-anchored membrane components produced by all mycobacteria (1, 9).

PIM biosynthesis begins on the inner leaflet of the plasma membrane where the mannosyltransferase PimA transfers a mannose from GDP-mannose to the inositol of the phospholipid phosphatidylinositol (PI) to produce PIM1 (**Fig. 1a**) (10). In a similar fashion, a second mannosyltransferase, PimB’, transfers the second mannose to the inositol of PIM1 to produce PIM2 (11, 12). PIM2 is then acylated on one of the mannose residues, and further mannosylated by unknown transferases to produce a tetra-mannosylated glycolipid (AcPIM4). The downstream fate of AcPIM4 is forked: AcPIM4 may either be mannosylated by the α1-2 mannosyltransferase PimE and another uncharacterized mannosyltransferase to form AcPIM6 (13), or AcPIM4 is further mannosylated by α1-6 mannosyltransferases such as MptA to produce LM, a lipoglycan similar to PIMs but with an elongated α1-6 mannose backbone (14, 15). Decorating the long mannan backbone of LM are multiple α1-2 linked mannoses added by the mannosyltransferase MptC (16–18). Finally, a branched arabinan domain may be added to the α1-6 mannan backbone to produce LAM. The biosynthesis of the arabinan domain is carried out by a suite of arabinosyltransferases including EmbC, as well as some of the Aft transferases though a detailed understanding of the involvement of each arabinosyltransferase is lacking (19–23).

**Figure 1.**
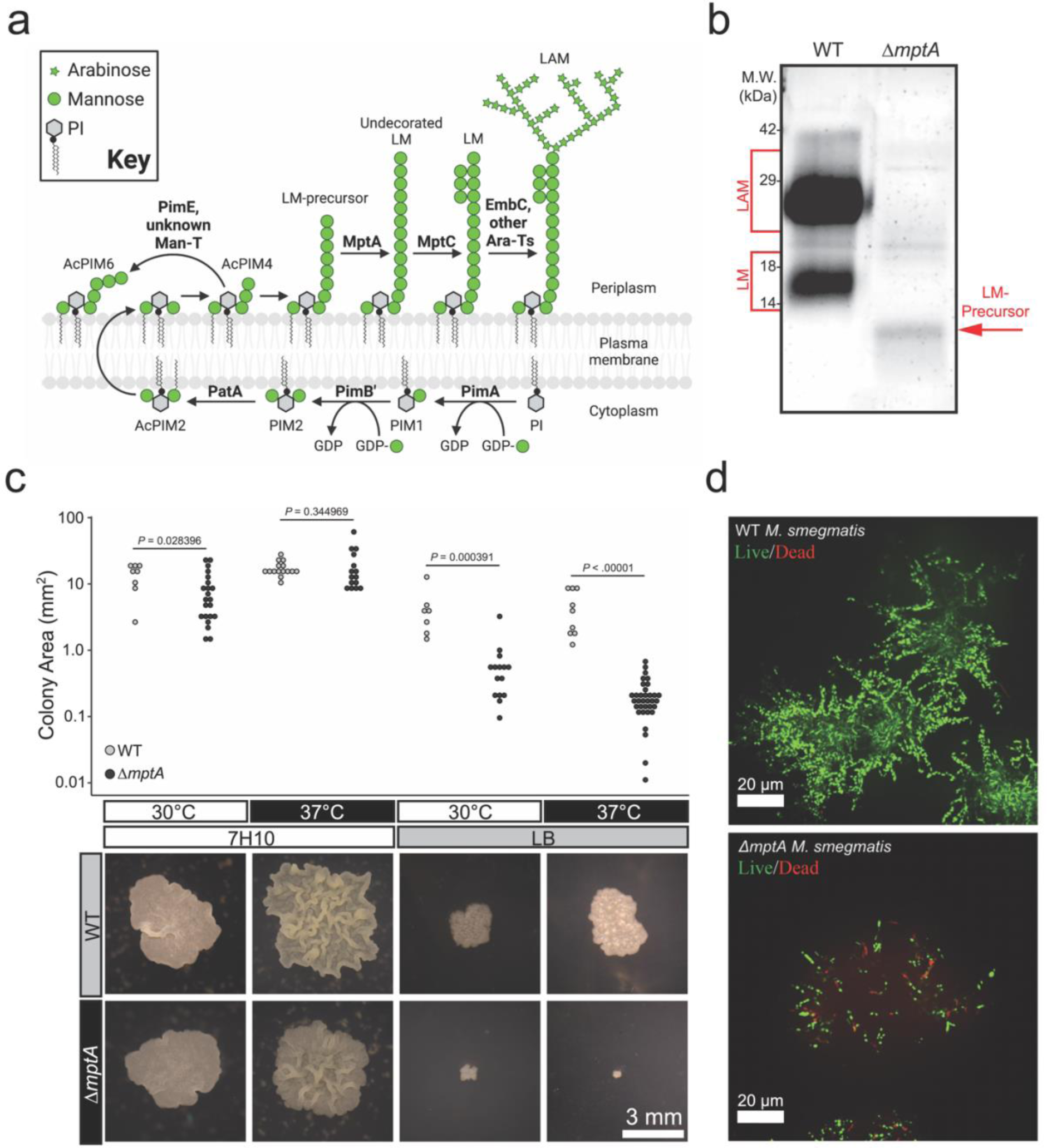
Medium-dependent growth defects of Δ*mptA*. **a)** PIM, LM, and LAM biosynthesis pathway. Bold text, enzymes. **b)** Lipoglycans extracted from WT and Δ*mptA* cells, visualized by glycan staining. **c)** Comparison of colony size between WT and Δ*mptA* on Middlebrook 7H10 and LB solid agar media grown at 30°C or 37°C. Top: quantification of colony area for each condition; bottom: example images of WT and Δ*mptA* colonies grown in each condition. **d)** Cross-sectional view of live/dead imaging of microcolonies from WT and Δ*mptA* strains stained with SYTO9 (live) and propidium iodide (dead) grown as micro-aggregates in LB medium.

PIMs, LM, and LAM specifically bind host receptors during infection to initiate both pro- and anti-inflammatory responses, as well as to prevent phagosome maturation, ultimately promoting intracellular survival (see review (24, 25)). While PIMs, LM and LAM are important virulence factors during mycobacterial infection, they are widely conserved among all mycobacteria, including many non-pathogenic species, suggesting a more fundamental physiological role in the cell envelope (26). This is supported by the essentiality of *pimA* and *pimB’* in nonpathogenic *Mycobacterium smegmatis* (10, 11). Furthermore, deficiencies in LM and LAM biosynthesis have demonstrated loss of fitness phenotypes in axenic culture. For example, overexpression of *mptC* in *M. smegmatis* and *M. tuberculosis* led to the truncation of the arabinan and mannan domains of LM and LAM and increased sensitivity to cell wall-targeting antibiotics (18, 27). An *M. smegmatis mptA* deletion mutant producing a small LM-precursor but no mature LM or LAM grew slowly on solid LB agar medium and failed to grow at 42°C (15). These observations reveal that LM and LAM may play an important role in mycobacterial growth and cell envelope integrity. Whether the phenotypes described in the literature share an etiological mechanism and what that mechanism might be remains unknown.

The goal of this study is to determine the physiological functions of mycobacterial lipoglycans. We generated various mutants of *M. smegmatis* that are deficient in LM and/or LAM biosynthesis to demonstrate that the arabinan domain is the key structural component of mycobacterial lipoglycans that define the subcellularly localized role for LAM in mycobacterial envelope maintenance and cell division.

## Results

### Defective colony growth of Δ*mptA* is dependent on culture media

An *mptA* deletion strain of *M. smegmatis* has a growth defect when grown on solid LB medium at 37°C (15). This contrasts with our previous observation that an *mptA* knockdown strain grows at a rate comparable to wildtype (WT) in Middlebrook 7H9 broth medium at 30°C (27), and suggests a conditional importance of LM and LAM for mycobacterial survival and growth. To better understand the role of MptA and its products LM and LAM, we generated Δ*mptA* and confirmed that it did not produce LM or LAM, but instead accumulated a small LM intermediate as reported previously **(Fig. 1b)**. We then grew Δ*mptA* on two different agar plates, LB or Middlebrook 7H10. As previously observed (15), Δ*mptA* grown on LB agar formed colonies significantly smaller than the WT grown either at 30 or 37°C **(Fig. 1c)**. In contrast, Δ*mptA* formed colonies that are comparable in size to the WT colonies when grown on Middlebrook 7H10 at either temperature **(Fig. 1c)**. These observations suggested that the growth defect of Δ*mptA* is medium-dependent. To examine the growth defect further, we conducted live/dead staining of *M. smegmatis* micro-colonies grown in LB in a glass-bottomed 96-well plate. WT and Δ*mptA* strains were stained with SYTO9 and propidium iodide such that live cells stain green and dead cells stain red. We found that WT micro-colonies had very few propidium iodide-positive dead cells **(Fig. 1d)**. In contrast, Δ*mptA* micro-colonies revealed many propidium iodide-stained cells, indicating Δ*mptA* is more prone to cell death when grown as micro-colonies in LB broth **(Fig. 1d)**. Collectively these results suggest that MptA and its biosynthetic products LM and LAM contribute to cell survival during aggregated growth conditions where LB was used as the medium.

### Δ*mptA* cannot maintain cell shape and lyses in pellicle growth

Next, we examined pellicle growth as it represents an established and scalable aggregated (biofilm) growth model, experimentally more tractable than colony growth. We compared growth of the WT strain and Δ*mptA* in M63 broth as we found that LB does not support robust pellicle growth of *M. smegmatis* (28–31). Because we observed cell death in LB-grown micro-colonies, we examined if Δ*mptA* similarly lyses in pellicle growth. In addition to Δ*mptA*, we used a previously established tetracycline-inducible conditional *mptA* knockdown strain (27). The pellicle of Δ*mptA* appeared comparable to that of the WT **(Fig. 2a)**. However, both Δ*mptA* and *mptA* knockdown cells induced by anhydrotetracycline (ATC) accumulated significantly higher levels of proteins in their culture medium than the WT (**Fig. 2b**). Mpa is a cytoplasmic protein, which is normally barely detectable in the spent medium. However, when *mptA* was knocked down by the addition of ATC, Mpa was readily detectable in the culture supernatant (**Fig. 2c**). Similarly, glycans and nucleic acid were released into the culture medium upon ATC-induced *mptA* knockdown **(Fig. 2d-e)**. Consistent with potential cell lysis, microscopic examination revealed morphological defects of Δ*mptA* and *mptA* knockdown (+ATC) cells **(Fig. 3a and d)**. We quantified morphological defects by measuring the cell width profile across the normalized cell length **(Fig. 3b)** and determining the distribution of maximum cell widths for each strain **(Fig. 3c)**. These quantitative analyses revealed that Δ*mptA* cells showed statistically significant morphological defects. Strikingly, the morphological defects were suppressed by growing the pellicle in osmo-protective M63 **(Fig. 3d-e)**, suggesting that cell shape deformation is dependent on the osmolarity of the medium. These data suggest that the turgor pressure is the force that deforms *mptA* knockdown cells and that the load-bearing function of the cell wall is possibly compromised in the absence of LM and LAM.

**Figure 2.**
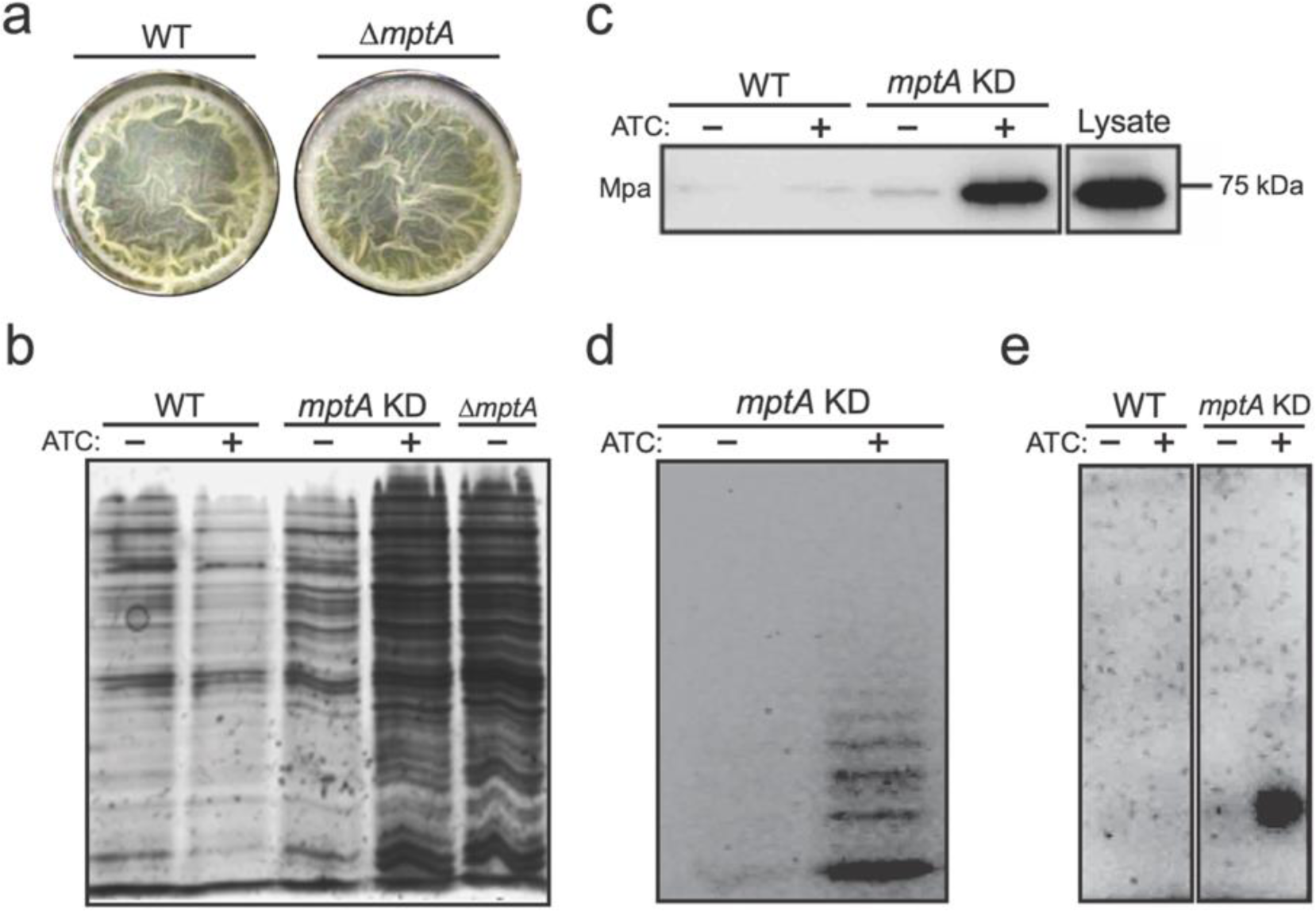
MptA-deficient cells lyse. **a)** Top-down view of WT and Δ*mptA* pellicle biofilms after 5-day growth in M63 at 37°C. **b)** SDS-PAGE of culture filtrates from WT, *mptA* KD, and Δ*mptA* pellicles visualized by silver staining. **c)** Western blot of WT and *mptA* KD pellicle culture filtrates to visualize the cytoplasmic marker protein Mpa. Lysate was diluted to match the protein concentration of the induced *mptA* KD pellicle culture filtrate as measured by absorbance at 280 nm. **d)** Fluorophore-assisted carbohydrate electrophoresis of the *mptA* KD pellicle culture filtrate. **e)** Agarose gel electrophoresis of nucleic acids in the pellicle culture filtrate visualized by ethidium bromide staining.

**Figure 3.**
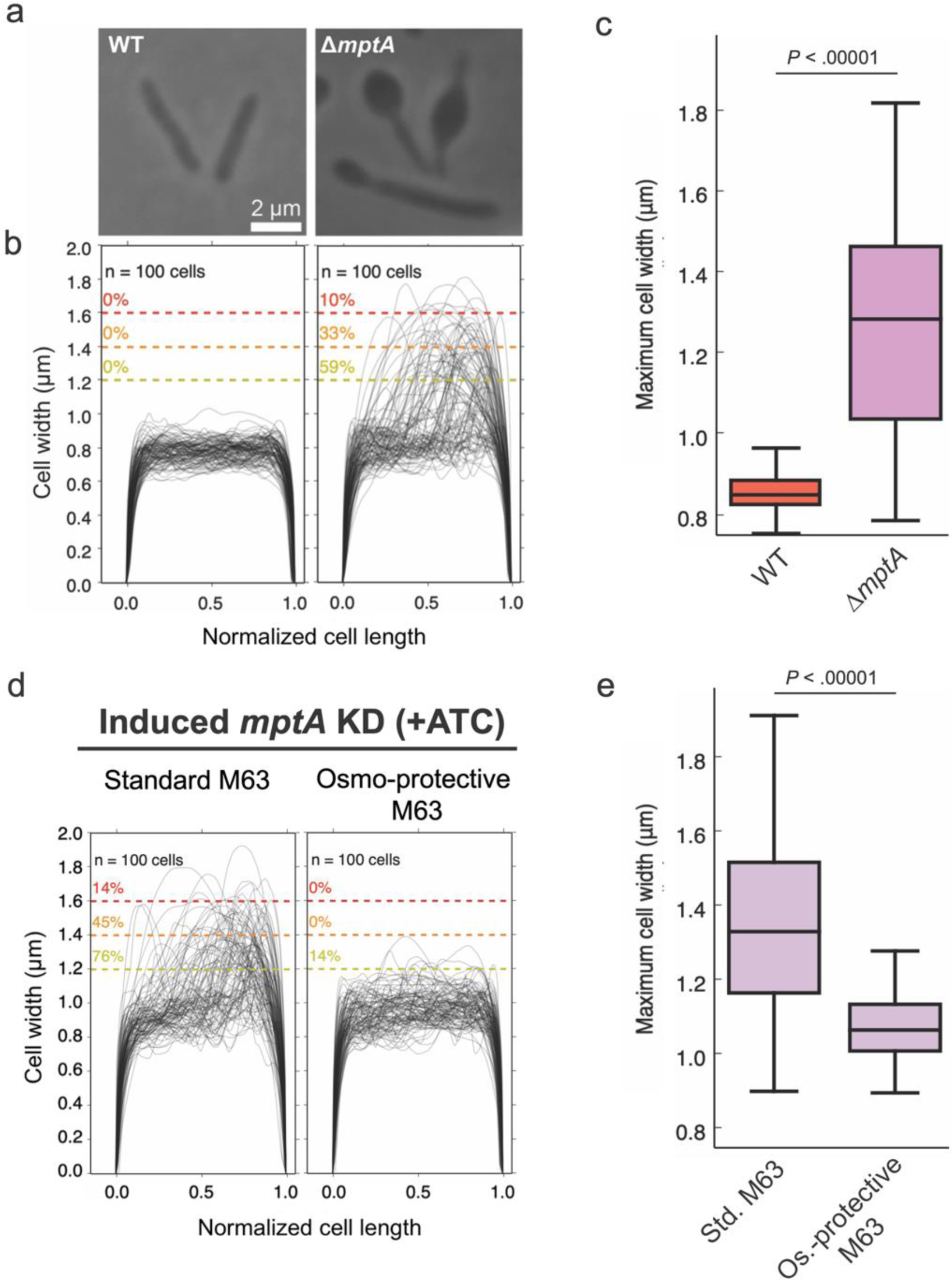
Defective cell morphology of Δ*mptA* grown as pellicle in M63 medium. **a)** Example phase micrographs of WT and Δ*mptA* cells grown as pellicles in M63 medium. **b)** Cell width profiles of both strains, with each cell’s length normalized to 1 (0.5 corresponds to midcell). The percentage values above the dotted colored lines indicate the portion of cells exhibiting maximum cell widths greater than or equal to the corresponding cell width threshold. **c)** boxplot comparing the distribution of maximum cell widths between WT and Δ*mptA* strains grown as pellicle biofilms. **d)** Cell width profiles of *mptA* KD cells induced with ATC grown as pellicle biofilms in standard (non-osmoprotective) M63 and osmoprotective M63 media. **e)** boxplot comparing the distribution of maximum cell widths between *mptA* KD pellicle biofilm cells grown in standard vs. osmoprotective M63 medium.

### Medium-dependent defective growth of Δ*mptA* is recapitulated in planktonic culture

The results so far suggested that the mutants struggle to maintain cell shape and lyse in aggregated growth. Nevertheless, our previous study showed that *mptA* knockdown cells display no defect when growing planktonically in Middlebrook 7H9 (27). We wondered if planktonically grown cells also show similar morphological defects in a medium-dependent manner. When we grew Δ*mptA* in Middlebrook 7H9 broth, they grew similarly to the WT as previously observed for *mptA* knockdown cells **(Fig. 4a)**. The cell morphology was normal **(Fig. 4b-c)**. In contrast, when Δ*mptA* was grown in LB broth, there was a growth defect, and its morphology was aberrant as observed in pellicle growth **(Fig. 4d-f)**. These data together suggest that the growth of Δ*mptA* is conditionally defective in a medium-dependent manner and is not specific to aggregated growth.

**Figure 4.**
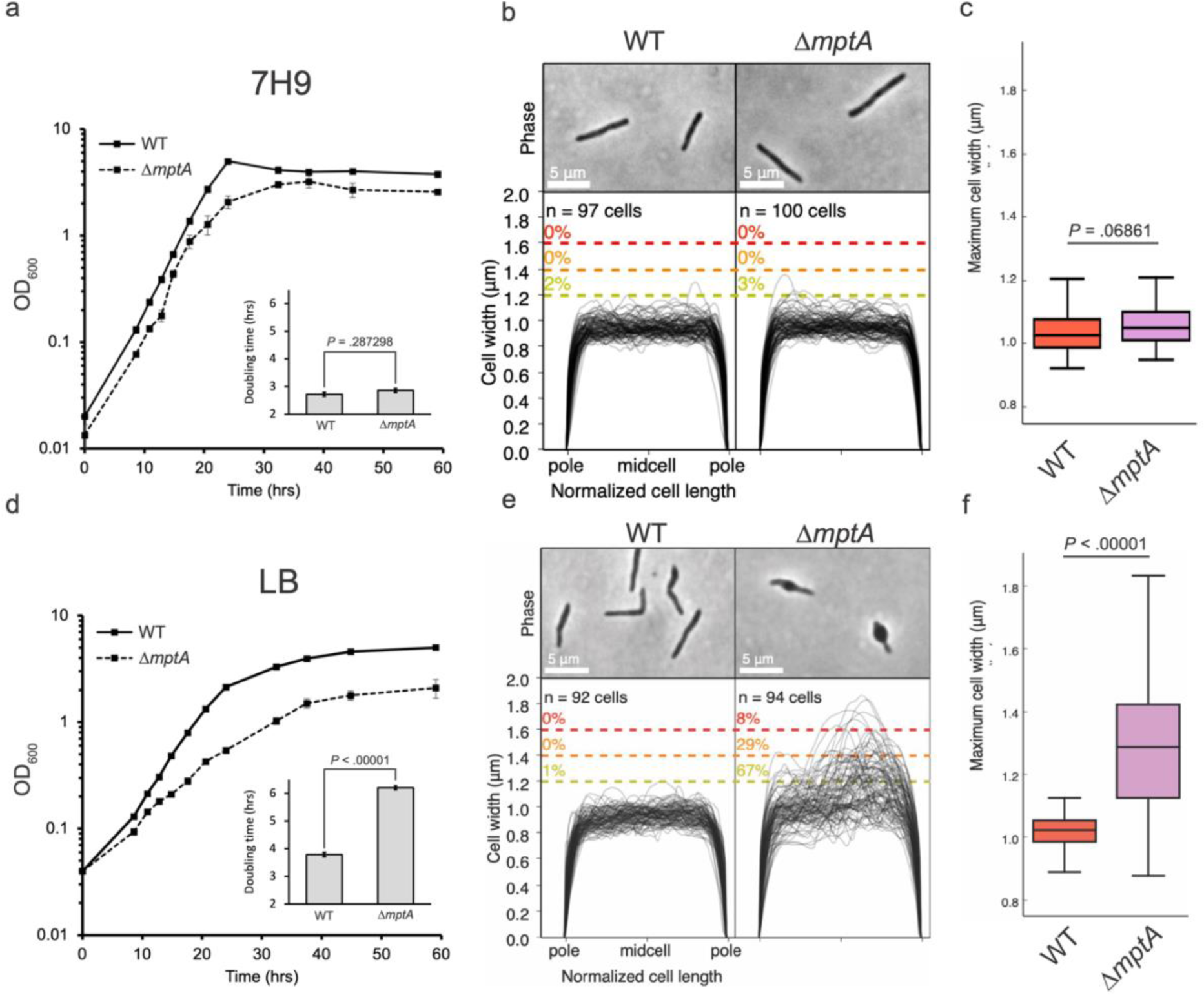
Growth and cell morphology of planktonically growing cells. **a and d**) growth curves of WT and Δ*mptA* cells grown planktonically in Middlebrook 7H9 (a) or LB (d) medium. Insets: doubling times of WT and Δ*mptA* in 7H9 (a, inset) or LB (d, inset) during log-phase growth. **b and e)** phase micrographs and cell width profiles of planktonic WT and Δ*mptA* cells in 7H9 (b) and LB (e). **c and f)** boxplots comparing the distribution of maximum cell widths between WT and Δ*mptA* strains grown planktonically in 7H9 (c) or LB (f).

### Δ*mptA* is hypersensitive to beta-lactam antibiotics

The suppression of the morphological defect when *mptA* KD cells were grown in high-osmolarity growth medium (see **Fig. 3d**) suggests that the turgor pressure is the force responsible for deforming *mptA* KD cell envelope and that the load-bearing function of the cell wall is compromised in the absence of LM and LAM. Previous work showed that *mptC* overexpression (OE), which stunts lipoglycan size, led to increased sensitivity to cell envelope-targeting antibiotics (18, 27), further suggesting that diminutive lipoglycans may compromise envelope integrity. Since Δ*mptA* produces even less developed lipoglycans than *mptC* OE, we tested whether this strain is similarly sensitive to cell envelope-targeting antibiotics. We determined the MIC of 5 different drugs for WT and Δ*mptA M. smegmatis*. We found that Δ*mptA* is over 40 times more sensitive to ampicillin+sulbactam and 8 times more sensitive to meropenem+sulbactam than WT (**Table 1**). Increased sensitivity of Δ*mptA* to the non-beta-lactam drugs was minor in comparison, indicating a specific sensitivity towards drugs targeting peptidoglycan crosslinking in cells deficient in LM and LAM.

**Table 1.** Δ*mptA* is more sensitive to β-lactam antibiotics. MIC, minimum inhibitory concentration. A non-growth-inhibitory dose of 50 µg/mL sulbactam was added to inhibit β-lactamases. Known primary drug targets are: isoniazid, the outer membrane biosynthesis; vancomycin and moenomycin, cell wall transglycosylation; ampicillin, cell wall transpeptidation (primarily D-D crosslinks); meropenem, cell wall transpeptidation (primarily L-D crosslinks); D-cycloserine, cytoplasmic cell wall precursor synthesis; and kanamycin, ribosome.

| Antibiotic<br>(+ 50 µg/mL sulbactam) | MIC (µg/mL) |  | Fold change in sensitivity<br>(*ΔmptA* /WT) |
| --- | --- | --- | --- |
|  | WT | *ΔmptA* |  |
| Isoniazid | 25.0 | 20.8 | 1.2 |
| Vancomycin | 0.8 | 0.4 | 2.0 |
| Moenomycin | 0.8 | 0.2 | 4.0 |
| Ampicillin | 8.3 | 0.2 | 42.7 |
| Meropenem | 6.3 | 0.8 | 8.0 |
| D-cycloserine | 83.3 | 50.0 | 1.7 |
| Kanamycin | 2.1 | 0.8 | 2.7 |

### LAM-deficient cells fail to maintain cell shape

Δ*mptA*’s increased sensitivity to beta-lactam antibiotics suggests that the absence of LM and LAM weakens the structural integrity of the peptidoglycan cell wall. Such defects in mycobacteria are commonly associated with characteristic “blebbing” of the cell, indicating that the cell wall is no longer strong enough to maintain cell shape against the turgor pressure (32–34), which is in fact what we observed for the Δ*mptA* and *mptA* KD strains **(see Figs. 3 and 4)**. Utilizing a collection of other *M. smegmatis* lipoglycan mutants, we set out to determine which specific structural component of LM/LAM lost in the Δ*mptA* strain is essential for maintaining cell shape. We confirmed the expected lipoglycan profiles of each strain under our experimental conditions, planktonic LB culture, by harvesting log phase cells, extracting their LM and LAM, and visualizing LM and LAM by SDS-PAGE **(Fig. 5a)**. We then analyzed the cell width of two strains that produce LAM but do not accumulate LM, Δ*mptC* and Δ*mptA L5::mptA-dendra2-flag*. Δ*mptA L5::mptA-dendra2-flag* was constructed by integrating a constitutive expression vector expressing a fluorescent Dendra2-FLAG-tagged MptA fusion protein into the L5 *attB* site on the chromosome of Δ*mptA.* This was done in an attempt to complement the Δ*mptA* strain, however, LAM biosynthesis was restored while LM did not accumulate, indicating partial complementation. Δ*mptC* and Δ*mptA L5::mptA-dendra2-flag* cells maintained rod shape similar to WT **(Fig. 5b-c)**. These results indicate that LM is not critical for maintaining envelope integrity and suggest that LAM is the important lipoglycan for envelope integrity. To further investigate, we analyzed the cell width of an MptC overexpression strain previously shown to produce LM and LAM with both dwarfed mannan and arabinan domains (18). We reasoned that this strain should display an intermediate morphology defect since it produces a LAM molecule with a small arabinan domain (18). As expected, the *mptC* OE strain had a significant blebbing defect, but with only about half the percentage of blebbed cells as were observed in Δ*mptA* **(Fig. 5b-c)**. These results suggest that the presence and correct size of the arabinan domain are both required for optimal cell wall integrity. To test this more directly, we knocked down the expression of the gene encoding the key arabinosyltransferase EmbC by ATC-inducible CRISPRi gene knockdown. When this strain was grown planktonically in LB with ATC, a subset of cells displayed a blebbed morphology like that seen for Δ*mptA* and *mptC* OE strains, further demonstrating the importance of the arabinan domain of LAM in maintaining cell shape **(Fig. 5b-c)**.

**Figure 5.**
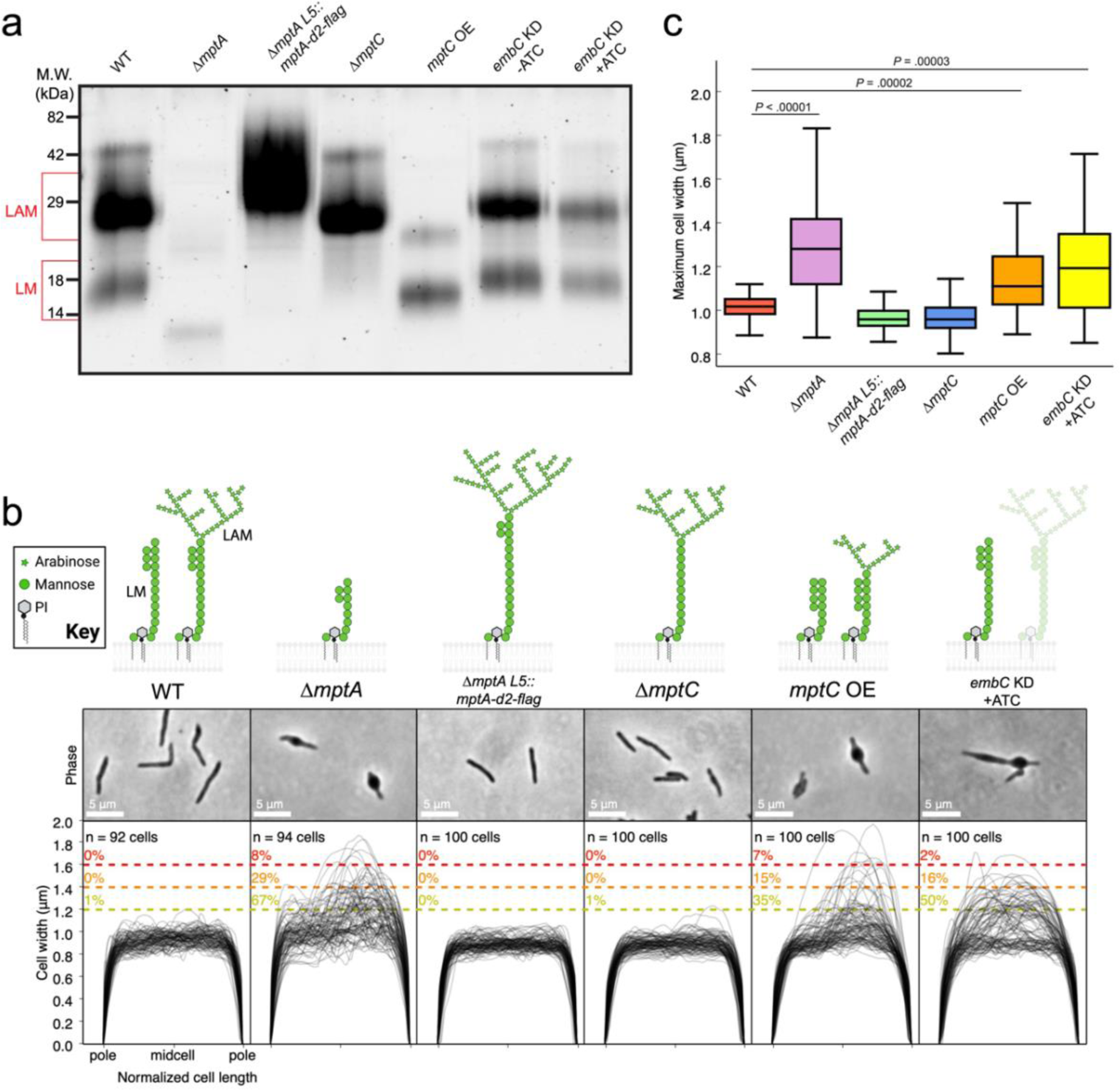
Cells deficient in LAM fail to maintain cell shape. **a)** SDS-PAGE of lipoglycans extracted from WT and various lipoglycan mutants, visualized by glycan staining. **b)** top: cartoon of lipoglycans for each strain. Bottom: phase micrographs and cell width profiles for each strain when grown planktonically in LB medium. **c)** Boxplots comparing the distribution of maximum cell widths between each strain.

### Cell envelope deformations in LM/LAM-deficient cells are associated with new poles and exhibit increased peptidoglycan remodeling

We noticed that many of the cell wall deformations were polar, potentially indicating that the cell wall defect is associated with elongation or division. Due to mycobacteria’s asymmetric growth, we can determine whether the blebs are associated with the old pole (cell elongation) or the new pole (septation and division). To determine cell polarity, we incubated cells with a fluorescent D-amino acid (FDAA) analog, called RADA, which are actively incorporated into growing cell walls by endogenous cross-linking enzymes and have been used extensively to study the remodeling of peptidoglycan cell walls in mycobacteria (35). As expected, WT and Δ*mptA* cells grown in LB planktonic culture were labeled asymmetrically at the cell poles, with the old pole showing brighter labeling extending farther into the sidewall than the new pole. We then aligned each non-septated blebbed cell according to old/new pole determined by its RADA labeling and plotted the cell width profiles. The majority of maximum cell widths for non-septated blebbed cells associated with new poles, though some blebs were associated with the midcell **(Fig. 6a)**. Intriguingly, Δ*mptA* cells also displayed higher RADA labeling at the sidewall, particularly at deformed regions of the cell envelope on the new pole half of the cell, indicating increased peptidoglycan remodeling along the sections of the cell envelope that are not actively growing **(Fig. 6b-c)**. This increased FDAA labeling at the regions of envelope deformations suggests that intense remodeling was induced to repair damaged or weak peptidoglycan cell wall. These data suggest that LAM is specifically important for peptidoglycan integrity during septation and daughter cell separation and not elongation.

**Figure 6.**
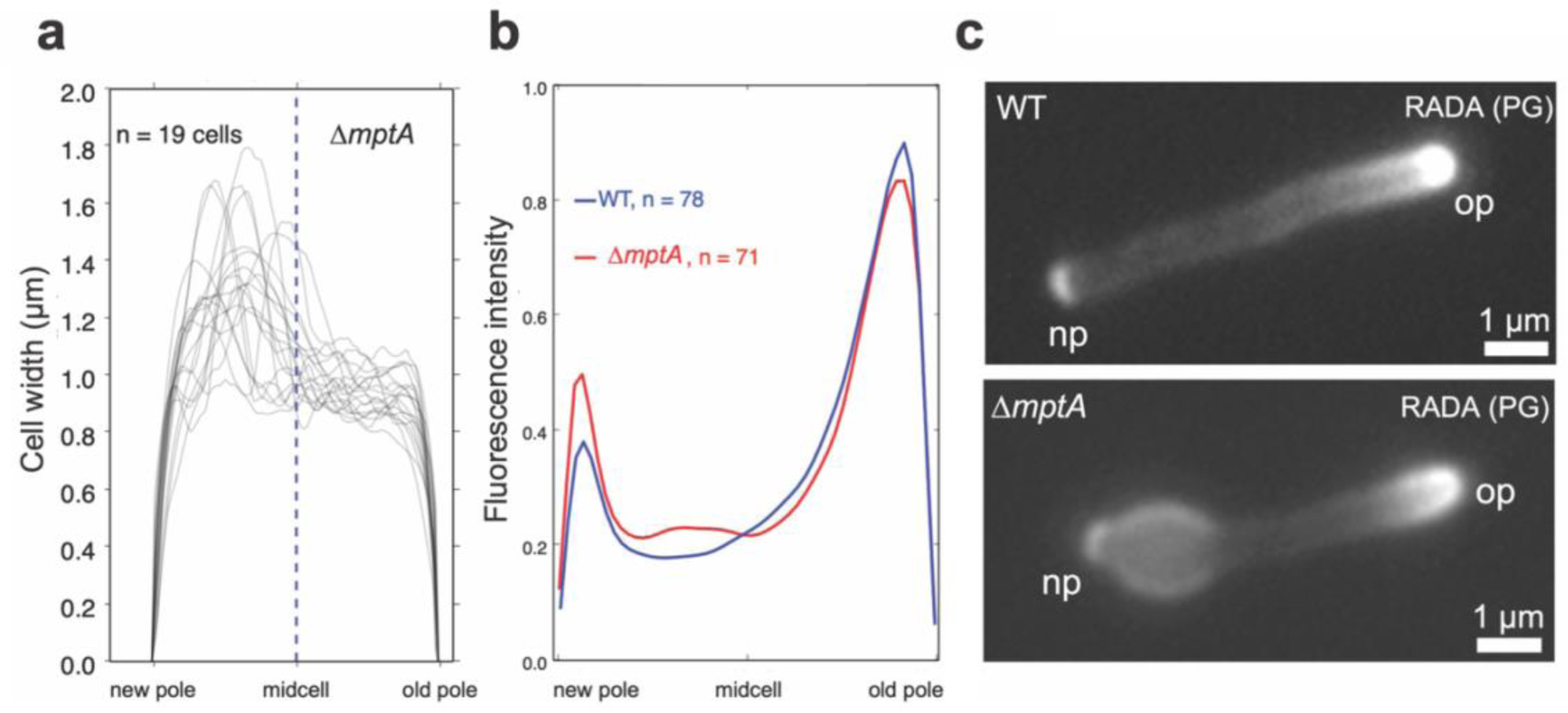
Cell envelope defects associate with the new pole. **a)** New-pole-aligned cell width profiles of deformed, non-septated Δ*mptA* cells grown planktonically in LB medium. Cells were labelled with RADA and aligned new pole (bright) to old pole (dim) based on RADA labelling. **b)** New-pole-aligned RADA labelling profile of WT and Δ*mptA* cells grown planktonically in LB medium. **c)** Example fluorescence microscopy images of RADA-labelled WT and Δ*mptA* cells. np, the new pole; op, the old pole; PG, peptidoglycan.

### MptA-Dendra2-FLAG localizes to cell septa

Taking advantage of the fluorescent protein tagged MptA in our Δ*mptA L5::mptA-dendra2-flag* we sought to determine the subcellular localization of MptA by fluorescent microscopy. In actively growing log-phase culture, MptA-Dendra2-FLAG accumulated as strong puncta, often in the middle of the cell or at a pole **(Fig. 7a)**. To determine whether the mid-cell signals are associated with septa, cells were incubated with HADA, another FDAA, to stain for peptidoglycan. MptA-Dendra2-FLAG localized with some but not all septa, supporting a role for MptA in septal biosynthesis or daughter cell separation **(Fig. 7b)**.

**Figure 7.**
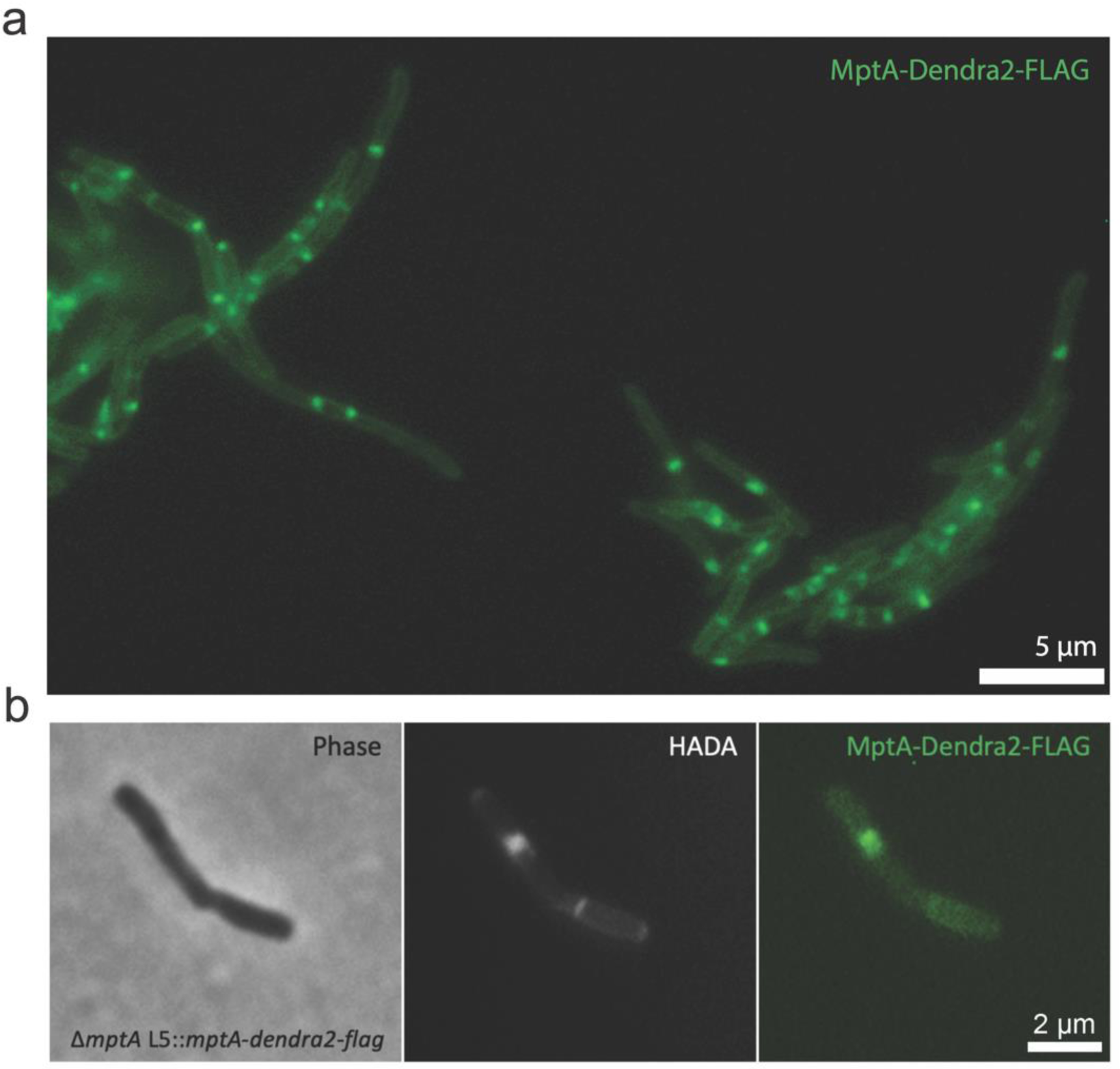
MptA-Dendra2-FLAG localizes to the septum. **a)** Example fluorescence microscopy image of MptA-Dendra2-FLAG expressed in Δ*mptA* background. **b)** Colocalization of HADA stained septum with MptA-Dendra2-FLAG.

### Expression of the cell wall synthase PonA1 restores envelope integrity in LM/LAM-deficient cells via a RipAB-independent mechanism

The new pole/septal location of cell envelope deformation suggests that LAM may function as a regulator of septal peptidoglycan hydrolase activity, which is required for daughter cell separation but must be carefully balanced against cell wall biosynthesis and remodeling to avoid lysis. Septal hydrolases must be able to cut the peptidoglycan connecting two daughter cells without destroying the peptidoglycan associated directly with each daughter cell’s inner membrane. This implies that some periplasmic factor spatially restricts the activity of septal hydrolases to avoid indiscriminate cell wall damage around the septum/new pole. The predominant septal hydrolase in *M. smegmatis* is RipA, which is directly inhibited by the PBP PonA1 through a RipA-binding motif on PonA1’s C-terminus **(Fig. 8a)** (33, 36–38). To test whether RipA inhibition could prevent envelope blebbing, we constitutively overexpressed PonA1 in Δ*mptA*. Strikingly, Δ*mptA* L5::*ponA1* maintained rod shape cell morphology **(Fig. 8b-c)**, indicating that the expression of PonA1 successfully rescued the peptidoglycan defect in the LM/LAM deficient Δ*mptA* mutant. Since PonA1 also catalyzes the biosynthesis of new peptidoglycan, it is possible that *de novo* peptidoglycan synthesis by PonA1’s enzymatic activity is responsible for rescuing Δ*mptA*. To test this, we expressed catalytically inactivated variants of PonA1 in the Δ*mptA* genetic background strain and measured cell width profiles. Expression of transpeptidase deficient (TP-), transglycosylase deficient (TG-), or catalytically dead (TP/TG-) PonA1 also rescued Δ*mptA* cell morphology, but to a lesser extent than WT PonA1 **(Fig. 8b-c)**. These data suggest that PonA1 can partially rescue the cell wall defects of Δ*mptA* through a non-catalytic function, potentially through RipA inhibition. To directly determine whether the cell wall defect of Δ*mptA* is dependent on RipA and its downstream operon partner RipB, an ATC-inducible *ripAB* CRISPRi knockdown construct was integrated into the L5 site of the Δ*mptA* strain (Δ*mptA L5::ripAB* KD) and the WT strain (WT *L5::ripAB* KD). When ATC was not added to the culture medium, each uninduced strain mimicked the cell morphology phenotype of their parental strains: Δ*mptA L5::ripAB* KD blebbed and WT *L5::ripAB* KD formed healthy rod-shaped cells **(Fig. 8d)**. As expected, induction of *ripAB* knock down via addition of ATC resulted in cell elongation and ectopic pole formation/branching in both genetic backgrounds, indicative of failed division in the absence of the key septal hydrolase, while the cell width characteristics of each strain remained similar to uninduced conditions **(Fig. 8d)**. Notably, healthy cell width characteristics and peptidoglycan remodeling were not restored in Δ*mptA L5::ripAB* KD upon knock down of the key septal hydrolase RipA, indicating that the cell wall defect of Δ*mptA* is not due to the dysregulation of RipA and RipB.

**Figure 8.**
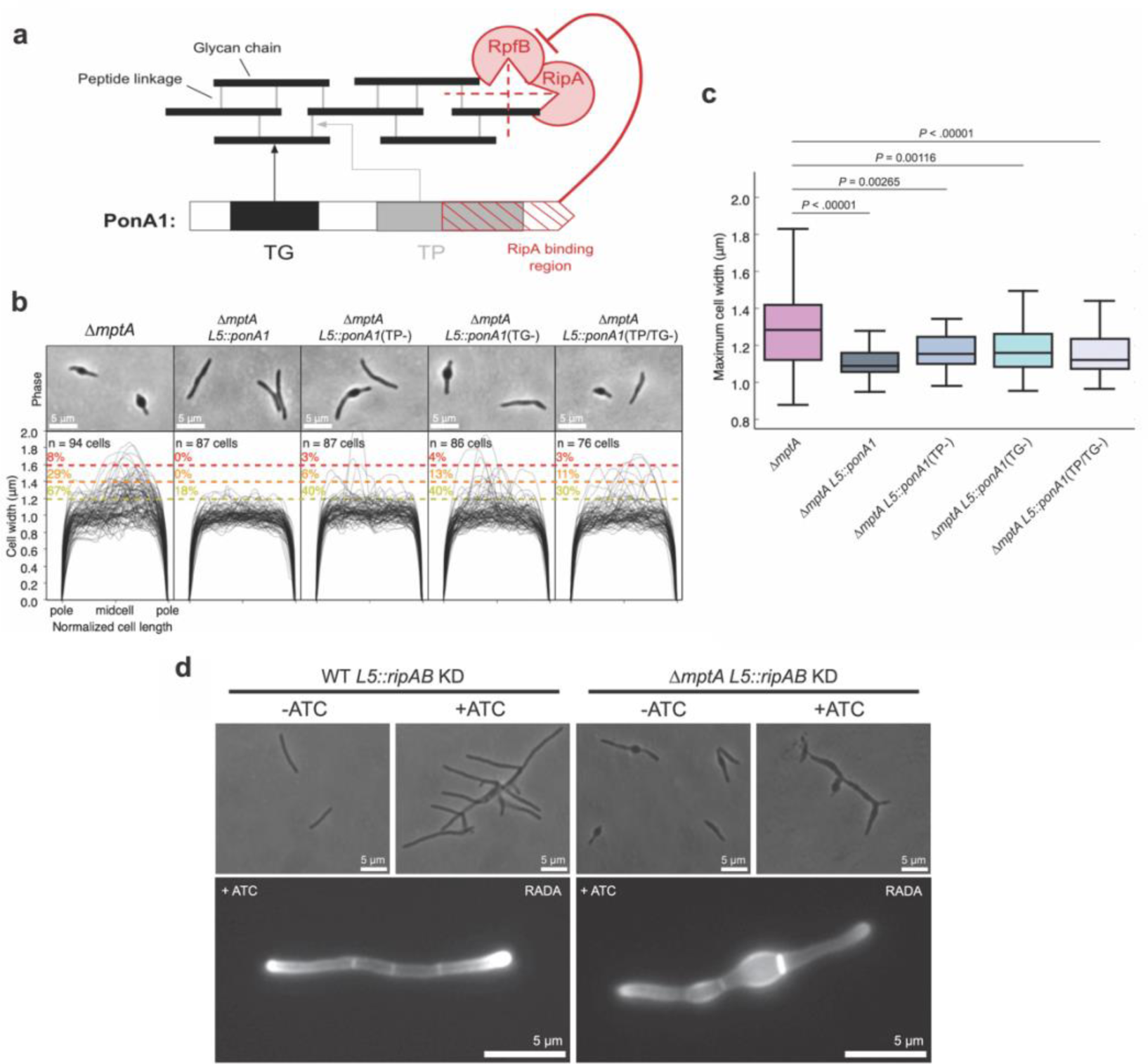
PonA1 rescues the morphological defects of Δ*mptA*. **a)** The domain structure of PonA1 and the corresponding cell wall substrates and interacting partners. TG, transglycosylase domain; TP, transpeptidase domain. The dotted red lines indicate which peptidoglycan structures are hydrolyzed by each component of the cell wall hydrolase complex comprised of RipA and RpfB. **b)** Phase micrographs and cell width profiles of Δ*mptA* cells overexpressing nothing (Δ*mptA*), fully functional PonA1 (Δ*mptA* + PonA1), transpeptidase-deficient PonA1 (Δ*mptA* + PonA1(TP-)), transglycosylase-deficient PonA1 (Δ*mptA* + PonA1(TG-)), and transpeptidase/transglycosylase deficient PonA1 (Δ*mptA* + PonA1(TP/TG-)). **c)** Boxplots comparing the distribution of maximum cell widths between Δ*mptA* and each PonA1 overexpression strain. **d)** (Top) Phase micrographs of ATC-inducible *ripAB* CRISPRi knock down strains constructed in the WT and Δ*mptA* genetic backgrounds. (Bottom) Fluorescent micrographs of ATC-inducible *ripAB* knock down strains stained with RADA.

### Cells producing a large LAM are defective in cell separation

One interesting consequence of complementing Δ*mptA* with the strong constitutive expression of *mptA-dendr2-flag* was that a larger LAM molecule was produced than in WT. Since dwarfed LAM weakens cell wall near the septum, a larger LAM may have the opposite effect of preventing the weakening of cell wall, like the *ripA*-deficient mutant, which cannot separate the two daughter cells and becomes multi-septated. To determine whether any potential impacts of a larger LAM molecule on the cell wall affect cell division, the Δ*mptA* L5::*mptA-dendra2-flag* strain was labeled with the FDAA peptidoglycan probe HADA, which indicated a hyperseptation phenotype in a subset of cells **(Fig. 9a)**. We quantified the percentage of septated and multiseptated cells in WT, Δ*mptA*, and Δ*mptA* L5::*mptA-dendra2-flag* to determine the prevalence of this cell division defect at the population level **(Fig. 9b)**. While only 20% of WT and Δ*mptA* cells were septated of which 2% were multiseptated, 48% of Δ*mptA* L5::*mptA-dendra2-flag* cells were septated of which 31% were multiseptated, demonstrating that strong constitutive expression of *mptA-dendra2-flag* has an inhibitory effect on septal resolution. Since LM does not accumulate in the Δ*mptA* L5::*mptA-dendra2-flag* strain, the division defect could either result from the lack of LM or the presence of larger LAM. To test this, we observed the level of septation in the Δ*mptC* strain, which also does not accumulate LM but does not produce a large LAM molecule **(see Fig. 5a-b)**. 24% of Δ*mptC* cells had a single septum and none were multiseptated, indicating that the lack of LM in this strain did not lead to a drastic increase in septation over WT. Furthermore, we observed hyperseptation in a strain overexpressing untagged *mptA* from an *hsp60* promoter on an episomal plasmid (*mptA* OE), a strain that was previously shown to produce LM and large LAM (18) **(Fig. 9a-b)**. The corresponding empty vector control strain exhibited WT levels of septation demonstrating that the effect was not due to the presence of the plasmid. Together, these results indicate that abnormally large LAM, not the lack of LM, causes the division defect.

**Figure 9.**
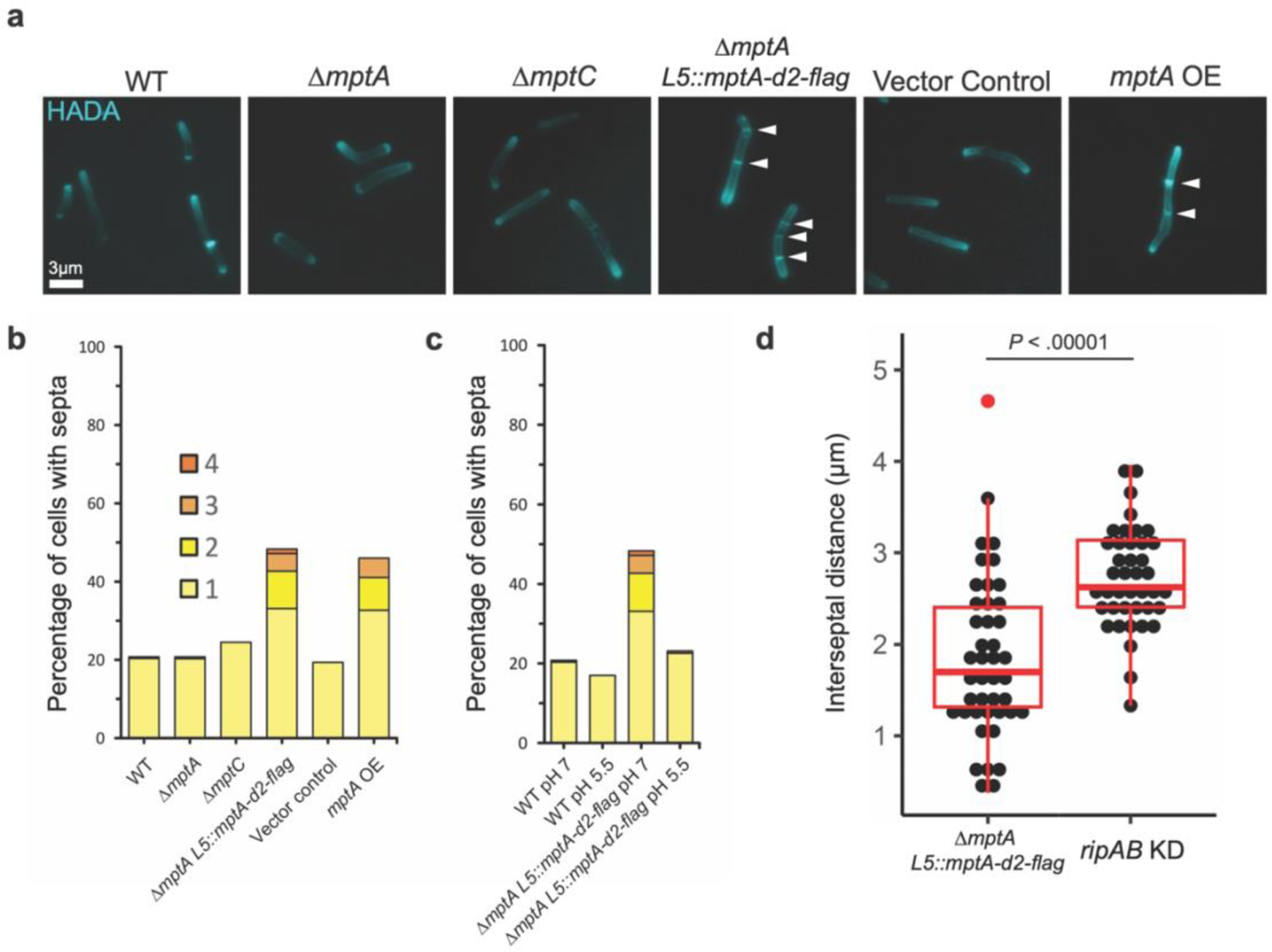
Large LAM results in multiseptated cells. **a)** Fluorescence micrographs of HADA-labelled cells. **b)** Septa enumeration for WT and lipoglycan mutant cells grown planktonically in LB. The combined bar height indicates the total percentage of cells with one or more septa. **c)** Septa enumeration for WT and Δ*mptA* L5::*mptA-dendra2-flag* (*mptA* comp) cells grown planktonically in neutral (pH 7) or acidic (pH 5.5) LB medium. **d)** Comparison between the distribution of septum-to-septum distances observed in multiseptated Δ*mptA* L5::*mptA-dendra2-flag* cells and multiseptated WT L5::*ripAB* KD (*ripAB* KD) cells.

A previous study found that in *M. tuberculosis,* the protease MarP proteolytically activates the septal peptidoglycan hydrolase RipA in response to low pH (39). MarP is conserved in *M. smegmatis*. We therefore reasoned that acidic growth conditions, which increase the activity of RipA, may increase septal hydrolysis enough to resolve the multiseptation effect of caused by oversized LAM in Δ*mptA* L5::*mptA-dendra2-flag*. To test this, we grew WT and Δ*mptA* L5::*mptA-dendra2-flag* in LB adjusted to pH 5.5 and compared the level of septation with that observed when cells were grown in standard LB (pH = 7) **(Fig. 9c)**. The acidic growth condition nullified the hyperseptation phenotype, suggesting that increased RipA activity may rescue this division defect. While increased RipA activity may help cells with large LAM to divide, LAM’s function does not appear to be directly involved in modulating RipA activity *per se* as we observed that knocking down this hydrolase did not rescue cells deficient in LAM (see **Fig. 8d**). If this is indeed the case, then the division defect caused by large LAM may be phenotypically distinct from that of cells deficient in RipA. In support of this, we found that while both Δ*mptA* L5::*mptA-dendra2-flag* and induced WT *L5::ripAB* KD have a multiseptation defect, the spacing between septa is markedly different between the two strains **(Fig. 9d)**. Knocking down *ripAB* results in consistent interseptal distances, indicative of regular cell wall elongation followed by failed cell separation, while Δ*mptA* L5::*mptA-dendra2-flag* displays much less consistent and shorter interseptal distances, indicative of either aberrant/premature placement of septa or a defect in cell elongation. Overall, these data support a model in which the size of LAM governs cell envelope structure at sites of division in such a way that impacts the success or failure of cell division.

## Discussion

In 1975, Norman Shaw proposed to use the term lipoglycan for Gram-positive macroamphiphiles that are structurally and functionally distinct from Gram-negative lipopolysaccharides (40). Lipoglycans are widely found in Gram-positive bacteria, including Actinobacteria, that do not produce lipoteichoic acids (LTAs). This intriguing observation led to a speculation that LTAs and lipoglycans fulfill functionally equivalent roles (41–43), but the analogy has been purely speculative. In the current study, we showed that the mycobacterial lipoglycan, LAM, regulates septation and division through modulating cell wall integrity. A similar role in septation and cell division has been ascribed to LTAs in several Gram-positive bacteria (44–47). In particular, *S. aureus* LTA biosynthesis proteins localize to division sites and interact directly with multiple divisome and cell wall synthesis proteins (48), and *S. aureus* mutants lacking LTA display aberrant septation and division as well as shape defects associated with cell lysis (46, 49). Furthermore, excessively long LTAs result in defective cell division, leading to cell “chaining” phenotypes (50). These phenotypes are strikingly similar to what we observed in *M. smegmatis* mutants, suggesting the universal importance of lipopolymer such as LTAs and LAM in governing cell division. Since LTAs and LAM are structurally different, we speculate that convergent evolution led these two distinct membrane-bound polymers to play a functionally similar role in controlling peptidoglycan dynamics.

Our results corroborate and unify previous findings suggesting the medium-dependent importance of LAM in mycobacterial growth and envelope integrity. The early report of Δ*mptA*’s small colony phenotype on LB was the first indication that LM and/or LAM were important for mycobacterial viability (15). In contrast, a later report showed that a *mptA* conditional knockdown strain grew normally in Middlebrook 7H9 growth medium (27). Our current study verified the cause of the contrasting observations in the previous studies and demonstrated the importance of culture medium in determining the fitness of Δ*mptA*. The role of LAM on growth is further supported by the observation of a similar small colony phenotype on complex solid medium for an *M. smegmatis lpqW* deletion mutant. LpqW acts at the branch point in PIM/LM/LAM biosynthetic pathway to shunt AcPIM4 more towards LM/LAM biosynthesis than AcPIM6 biosynthesis (16). Presumably due to the redirection of AcPIM4 to AcPIM6 synthesis, Δ*lpqW* failed to accumulate LM and LAM (16). The *lpqW* deletion strain was found to consistently accumulate mutations in *pimE*, preventing the conversion of AcPIM4 to AcPIM6, when grown on complex medium like LB but not defined Middlebrook medium, resulting in restored colony size and biosynthesis of LM and LAM (26). The medium dependency of the *lpqW* mutant phenotypes is consistent with phenotypes resulting from disruption of *mptA*.

One clue as to how LM and LAM may be important for growth came from the observation that *mptC* overexpression truncated LM and LAM and concomitantly increased sensitivity to cell-wall targeting antibiotics (27). This observation suggested that lipoglycan deficiency may compromise the mycobacterial cell wall in some way. In this study, we demonstrated that peptidoglycan remodeling, and its load-bearing and shape maintenance functions are compromised in the absence of LM and LAM. Furthermore, we demonstrated that LM/LAM-deficient cells are specifically sensitive to inhibition of cell wall crosslinking, and that cell shape defects and de-localized peptidoglycan remodeling are subcellularly localized to regions associated with recent division, and not elongation in LM/LAM-deficient cells. By testing the cell morphology characteristics of five distinct LM/LAM biosynthetic mutants, we determined that full sized LAM, and not LM, is important for maintaining envelope integrity. A recent study indicated that the α1-6 mannan backbone of LM/LAM carries 13-18 mannose residues (51). Using CHARMM-GUI Glycan Modeler, the chain length of 13 α1-6-linked mannoses is predicted to be 4.2 nm (52, 53). Given the predicted height of the periplasmic space being 14-17 nm (54), the mannan backbone part of LM/LAM is not sufficient to span the periplasmic space and reach the peptidoglycan layer. However, ∼23 residues of α1-5-linked arabinose, found as a branch in the arabinan part of LAM (51), can extend LAM by 9.8 nm, which makes it long enough to reach the peptidoglycan layer. Multiple branches of these arabinan chains increase the surface area/coverage of the lipoglycan structure precisely where the molecule may interact with the cell wall or cell wall-acting proteins.

Cells lacking sufficient cell wall hydrolase activity fail to fully divide, resulting in a multi-septation phenotype (38). We initially hypothesized that LAM could be modulating septal hydrolase activity such that in LAM’s absence, hydrolase activity becomes too active and when LAM is too big, hydrolase activity is impaired. However, we determined that the key septal hydrolase, RipA, was not responsible for the cell shape defect observed in LAM-deficient cells, as knocking down *ripAB* expression in the Δ*mptA* background did not relieve cell shape defects. The compounded phenotypes of *mptA* deletion and *ripAB* knock down demonstrate that the cell wall defects of LAM-deficient cells near septa and new poles are independent of RipA-mediated cell separation. Additionally, we showed that the multiseptation defect in large LAM-producing cells was qualitatively different from the multiseptation defect in *ripAB* knockdown cells. The less regular interseptal distances of the Δ*mptA* L5::*mptA-dendra2-flag* strain suggest that LAM may govern proper septal placement or cell elongation immediately after septal placement.

Intriguingly, overexpression of catalytically dead PonA1 partially rescued Δ*mptA* even though we showed that RipA is not the cause of cell shape defects in Δ*mptA*. Dead PonA1 overexpression may rescue Δ*mptA* through interactions with other proteins. Apart from RipA, PonA1 is known to physically interact with another divisome protein LamA, which is involved in maintaining division asymmetry and drug resistance (55). PonA1 also interacts with the potential scaffold protein MSMEG_1285 (Rv0613c), which is known physically interact with additional cell envelope proteins (56). Since PonA1 is involved in both division and elongation, which are processes likely driven by large multi-protein divisome and elongasome complexes, PonA1 is likely to interact with additional proteins involved in growth and division (see reviews (57, 58)). These additional interactions may explain the RipA-independent partial rescue of Δ*mptA* with catalytically dead PonA1. Precise molecular mechanisms by which LAM governs the functions of mycobacterial divisome in a RipA-independent manner is an important topic of future research.

## Methods

### Growth of *M. smegmatis* mc^2^155

For planktonic growth, cells were grown in Middlebrook 7H9 supplemented with 15 mM NaCl, 0.2% (w/v) glucose, and 0.05% (v/v) Tween-80 or LB supplemented with 0.05% (v/v) Tween-80 and incubated at 37°C and 130 rpm until mid-log phase (OD600 = 0.5 to 1.0). For pellicle biofilm growth, 20 µL planktonic stationary phase culture was diluted 1:100 into 2 mL M63 medium in a 12-well plate and incubated for 3 – 5 days at 37°C without shaking. Spent culture medium was filtered through a 0.22 µm filter to remove residual cells. For colony growth, serial dilutions of planktonic culture were spread evenly over LB or Middlebrook 7H10 (supplemented with 15 mM NaCl and 0.2% (w/v) glucose) solid agar medium and incubated at 30°C or 37°C for 3 – 4 days. Colonies were photographed using Amhersham ImageQuant 800 (Cytiva) and colony area was determined using ImageJ software (59).

### Construction of Δ*mptA* strain

Genomic regions upstream and downstream of *MSMEG_4241* (*mptA)* were amplified by PCR using primers (5’-GACAGGACTCTAGCCAAAGAACATCGGTCCGGTGTACG -3’ and 5’-CGGCTCGCCGTCGTGGCCTAGGTGTGGACTGTCGAGCC -3’ for the upstream region and 5’-TAGGCCACGACGGCGAGC -3’ and 5’-GCTGTCAAACCTGCCAACTTATCACGCTGGTGGAAGTGAT -3’ for the downstream region) and assembled via HiFi cloning into pCOM1 (60). This construct was electroporated into WT *M. smegmatis* and clones that had incorporated the vector into the genome via single homologous recombination event were selected for on Middlebrook 7H10 plates containing 100 µg/mL hygromycin. A “single cross-over” mutant was isolated and grown to an OD of 1 in the absence of hygromycin to allow a second recombination event to complete the gene deletion and excise the vector backbone. The “double cross-over” strain was selected for on Middlebrook 7H10 plates containing 5% sucrose. Candidate clones were confirmed to be sensitive to hygromycin and resistant to sucrose indicating the loss of the vector backbone. The absence of *MSMEG_4241* (*mptA)* was confirmed by PCR with primers (5’-AGTACCTGCGCGAACGTC -3’ and 5’-TGAGCAGTTCGAAGGTCAGG -3’) using the genomic DNA of the mutant as a template.

### Lipoglycan extraction and analysis

Pellets from 20 mL log phase planktonic cultures were harvested and delipidated with chloroform/methanol/water as previously described (27, 61). Delipidated pellets were resuspended and incubated in phenol/water (1:1) at 55°C for 2 hours to extract LM and LAM. The aqueous phase containing lipoglycan was washed with an equal volume of chloroform. The resulting aqueous phase was then concentrated by vacuum centrifugation and resuspended in water to obtain purified LM and LAM. LM/LAM samples were incubated with 0.1 mg/mL Proteinase K for 1 hour at 50°C to remove residual proteins and separated by SDS-PAGE (15% gel) with a constant 130 V voltage using a Bio-Rad system. Lipoglycans on the separated gel were then stained using the ProQ Emerald 488 glycan staining kit (Life Technologies) and visualized using Amhersham ImageQuant 800 (Cytiva) as previously described (13, 18).

### Live/Dead staining of microcolonies

For microcolony growth, cells from frozen stock were incubated for 48-72 hours in liquid LB media supplemented with 0.05% Tween-80 under shaken conditions at 37°C. The culture was diluted 100x in fresh LB media (without Tween-80) and deposited in the center of a glass-bottomed well in a 96-well plate (MatTek P96G-1.5-5-F). The culture was then grown at 37°C under static conditions for approximately 48 hours. The medium was replaced with fresh LB media supplemented with 0.1% SYTO9 and 0.1% propidium iodide (Invitrogen) and incubated for 1 hour. The microcolonies were imaged using a confocal spinning disk unit (Yokogawa CSU-W1), mounted on a Nikon Eclipse Ti2 microscope with a 100x silicone oil immersion objective (N.A. = 1.35), and captured by a Photometrics Prime BSI CMOS camera.

### Protein analysis by SDS-PAGE and western blotting

Culture filtrate samples were separated by SDS-PAGE (12% gel) and stained by silver staining. For western blotting, the gel was transferred to a polyvinylidene difluoride (PVDF) membrane (Bio-Rad) in a transfer buffer (25 mM Tris-HCl (pH 8.3), 192 mM glycine, 0.1% SDS (w/v)) at 14 V for overnight. The membrane was blocked in a blocking buffer (5% skim milk in PBS + 0.05% Tween 20 (PBST)) for 2 hours, probed with rabbit anti-Mpa IgG diluted 1:2000 in the blocking buffer for 8 hours at 4°C, and washed 10 minutes 3 times in PBST before being incubated with horseradish peroxidase-conjugated anti-rabbit IgG (GE Healthcare) diluted 1:2000 in the blocking buffer for 1 hour at room temperature. Finally, the membrane was washed 10 minutes 3 times in PBST and developed with homemade ECL reagent (2.24 mM luminol, 0.43 mM *p*-coumaric acid, and 0.0036% H2O2 in 100 mM Tris-HCl (pH 9.35)) and imaged the chemiluminescence using an ImageQuant LAS-4000 mini Imaging System (GE Healthcare).

### Fluorophore-assisted carbohydrate electrophoresis of biofilm culture filtrates

Filtrates of spent medium from pellicle biofilm culture were concentrated 10x under a nitrogen stream. An aqueous phase containing carbohydrates was recovered from 50 µL of concentrated filtrate vortexed with 150 µL water-saturated butanol. The remaining butanol phase was washed with 50 µL of water to repeat the extraction. The combined aqueous phase was mixed with 400 µL chilled (−20°C) 100% ethanol to precipitate the crude carbohydrates. The precipitated carbohydrate pellet was briefly dried under a nitrogen stream and resuspended in 10 µL of 0.15 M 8-aminonaphthalene-1,3,6-trisulfonic acid (ANTS) dissolved in 15% acetic acid and 10 µL of 1 M sodium cyanoborohydride dissolved in DMSO. The mixture was incubated for 30 hours at 37°C. The ANTS-labelled samples were dried by vacuum centrifugation, resuspended in 55 µL water, and separated on a 30% polyacrylamide gel in a chilled gel box running at a constant voltage of 190 V for 2.5 hours. Separated carbohydrates were visualized on a UV transilluminator.

### Agarose electrophoresis of nucleic acids

Pellicle culture filtrates were mixed with 6X loading dye and run on a 1% agarose gel at 100 V for 32 minutes. The gel was incubated with 0.01% ethidium bromide for 10 minutes and visualized on a UV transilluminator.

### Microscopy and quantification of cell morphology

Log phase cells were dispensed onto an agar pad (1% agar in water) on a slide glass and imaged at 1000x magnification (100x objective lens, N.A. = 1.30) with a Nikon Eclipse E600 microscope. Coordinates of cell outlines were obtained by analysis of phase contrast micrographs analyzed with Oufti (62). These coordinates were then processed using an original python script (https://gitfront.io/r/user-9868917/FFpPeaQAUXtj/Supplemental-code-Sparks-2023/) to extract cell width profiles and maximum cell width values for each cell. Cell width profiles of individual cells were compiled to obtain population level cell width profiles.

### Determination of MIC

In a 96-well plate, antibiotics were serially diluted 1:2 in wells containing Middlebrook 7H9 growth medium containing 50 µg/mL sulbactam and inoculated with *M. smegmatis* culture to an OD600 of 0.03. The plate was incubated for 24 hours at 37°C before the addition of resazurin to a final concentration of 0.0015%. The plate was incubated for an additional 8 hours at 37°C after which the MIC was determined spectroscopically at 570 and 600 nm, as described previously (63).

### Construction of *embC* and *ripAB* CRISPRi KD strains

ATC-inducible CRISPRi knockdown L5 integration vectors expressing guide RNAs targeting *embC* and *ripAB* were obtained from the Mycobacterial Systems Resource and electroporated into WT or Δ*mptA M. smegmatis*. Transformants were selected for on Middlebrook 7H10 plates containing 20 µg/mL kanamycin and knockdown was induced with 50 ng/mL ATC.

### RADA and HADA labelling and quantification

To label short-term peptidoglycan remodeling, RADA was added to log phase cells at a concentration of 10 µM and incubated for 15 minutes at 37°C. Cells were washed twice with LB and imaged by phase and fluorescence microscopy. To quantify RADA fluorescence, non-septated cells were first outlined in Oufti from phase contrast micrographs. Fluorescent profiles from corresponding fluorescent micrographs were mapped to each cell and oriented from dim pole to bright pole. The highest values for each cell’s fluorescent profile were normalized to 1 and a high order polynomial regression curve was fitted to the population-level data to produce an average fluorescence profile for each strain. To fully label cell walls and septa, HADA was added to log phase cells at a concentration of 500 µM and incubated for 1 hour at 37°C. Cells were washed twice with LB and imaged by phase and fluorescence microscopy. The number of septa per cell was manually counted. The distances between septa in multiseptated cells were measured using ImageJ.

### Construction of Δ*mptA* L5::*mptA-dendra2-flag* strain

An L5 integration *mptA-dendra2-flag* expression vector was obtained from the Mycobacterial Systems Resource and electroporated into the Δ*mptA* strain. Transformants were selected for on Middlebrook 7H10 plates containing 12.5 µg/mL apramycin and expression of the fluorescent fusion protein was confirmed by microscopy.

### Site-directed mutagenesis of PonA1 overexpression strains

The WT *ponA1* gene (*MSMEG_6900*) was PCR-amplified from *M. smegmatis* genomic DNA using two primers (5’-AAAAAAAA<u>CATATG</u>AATAACGAAGGGCGCCACTCC -3’ and 5’-TTTTTT<u>ATCGAT</u>TCACGGAGGCGGCGGG -3’), which contain NdeI and ClaI sites (underlined) respectively. The PCR product was digested with NdeI and ClaI, and ligated into pMUM261, which was digested with the same enzymes. pMUM261 is a variant of pMUM110, an integrative expression vector driven by the weak P766-8G promoter (64). Specifically, in pMUM261, the 5’-UTR up to the HindIII site was replaced with <u>AAGCTT</u>TTTGGTATCATGGGGACCGCAAAGAAGAGGGG<u>CATATG</u> to remove an NcoI site and introduce an NdeI site at the start codon, and additionally 54 bp fragment containing multiple restriction enzyme sites (HpaI-ClaI-AflII-NheI-NcoI) located at the downstream of the TetR38 gene was removed by digesting with NcoI and HpaI, blunt-ending, and self-ligating. Because the P766-8G-driven expression of the resultant pMUM303 plasmid was too low to rescue Δ*mptA*, we then swapped in the promoter region of pMUM100 (a variant of pMUM106 (64), but carrying the strong P750 promoter) by digesting both with NheI and HindIII and ligating the relevant fragments, resulting in pMUM314, a P750-deriven PonA1 expression vector. Site-directed mutagenesis was performed using Platinum SuperFi DNA polymerase (Invitrogen) according to kit instructions. The primers, 5’-GGCCGCCCAGGACCGTGACTTCTAC -3’ and 5’-CGGTCCTGGGCGGCCATCACCGC -3’, were used to mutagenize the catalytic residue of the transglycosylase domain (E193Q, resulting in pMUM315). The primers 5’-GACGGGTGCGGCGTTCAAGGTGTTCGC -3’ and 5’-CTTGAACGCCGCACCCGTCGGCAGGCC -3’ were used to mutagenize the catalytic residues of the transpeptidase domain (S468A/S469A, resulting in pMUM316 or both E193Q and S468A/S469A, resulting in pMUM317). These expression vectors were integrated into the mycobacteriophage L5 attachment (*attB*) site of Δ*mptA’s* genome.

## Acknowledgements

YSM and JY were supported by NIH (R21AI168791 and DP2GM146253, respectively). We thank Dr. Sloan Siegrist (University of Massachusetts Amherst) for sharing her cell wall labeling expertise and reagents, Dr. Keith Derbyshire (Wadsworth Center) for CRISPRi constructs, and Dr. Heran Darwin (New York University) for anti-Mpa antibody.

